# A light sheet fluorescence microscopy protocol for *Caenorhabditis elegans* larvae and adults

**DOI:** 10.1101/2022.08.05.503008

**Authors:** Jayson J. Smith, Isabel W. Kenny, Carsten Wolff, Rachel Cray, Abhishek Kumar, David R. Sherwood, David Q. Matus

**Author notes:** **Correspondence:** David R. Sherwood, David Q. Matus. These authors contributed equally to this work and share first authorship (Jayson J. Smith, Isabel W. Kenny). D.Q.M is a paid consultant of: Arcadia Science.

## Abstract

Light sheet fluorescence microscopy (LSFM) has become a method of choice for live imaging because of its fast acquisition and reduced photobleaching and phototoxicity. Despite the strengths and growing availability of LSFM systems, no generalized LSFM mounting protocol has been adapted for live imaging of post-embryonic stages of *C. elegans*. A major challenge has been to develop methods to limit animal movement using a mounting media that matches the refractive index of the optical system. Here, we describe a simple mounting and immobilization protocol using a refractive-index matched UV-curable hydrogel within fluorinated ethylene propylene (FEP) tubes for efficient and reliable imaging of larval and adult *C. elegans* stages.

## 1. INTRODUCTION

Light sheet fluorescence microscopy (LSFM) affords several advantages for live imaging of biological samples over standard epifluorescence or confocal microscopy. Whereas wide-field microscopy illuminates an entire specimen for imaging, LSFM achieves reduced phototoxicity, photobleaching, and background signal by restricting the proportion of the sample that is illuminated during acquisition. Relative to wide-field imaging, point-scanning confocal methods reduce out of focus sample illumination in the X-Y dimension by only exciting a single point in the sample at a time. To cover the whole region of interest the laser repeatedly sweeps across the sample and for each point scanned the entire Z depth is illuminated. Thus, out of focus photobleaching and phototoxicity occurs in the Z-dimension (Fischer et al., 2011). In contrast to a confocal point-scanning microscope where out of focus light is rejected by discarding unwanted emitted photons, LSFM systems generate a light sheet that selectively illuminates a narrow z-range of the sample in the desired focal plane at a given time (Fischer et al. 2011; Albert-Smet et al. 2019). This eliminates out of focus photobleaching and permits the collection of the entire fluorescence signal of a section of the sample at one time point, dramatically increasing acquisition speeds (Fischer et al. 2011). Another advantage of LSFM is the ability to acquire multi-view image data via multidirectional illumination, sample rotation, or a combination of both techniques (Huisken and Stainier 2009; Schmid and Huisken 2015). To overcome loss of resolution at increased tissue depths, many LSFMs are equipped with the ability to simultaneously image an individual sample from multiple sides, which can then be computationally deconvolved and reconstructed to render a single image of isotropic resolution. These technical advantages have made LSFM a popular imaging method for visualization of complex three-dimensional cells and tissues over developmental time (Keller et al. 2008; Liu et al. 2018).

Most LSFMs are equipped with two or more perpendicular illumination and detection objectives with the sample centered under or between the objectives. This unique orientation of objectives relative to the sample impedes the use of traditional flat microscopy slide mounts for the majority of LSFM systems. Samples for LSFMs are thus often embedded in a cylinder of low-melt agarose that hangs vertically between the objectives. In cases where the agarose is not dense enough to maintain its form, rigid fluorinated ethylene-propylene (FEP) tubes can be used to surround the agarose cylinder to stabilize and support the agar (Kaufmann et al. 2012; Girstmair et al. 2016; Steuwe et al. 2020). The refractive indices of low-melt agarose (1.33) and FEP tubes (1.34) are well matched to the refractive index of water (1.33) and this sample mounting method works well for many organisms.

The *C. elegans* embryo has been particularly helpful in advancing the use of LSFM. For example, *C. elegans* embryogenesis was used to demonstrate the enhanced spatiotemporal resolution that is achieved using lattice light-sheet microscopy (Chen et al. 2014). Similarly, the *C. elegans* embryo facilitated showing the effectiveness of four-dimensional (4D) live imaging with the Dual Inverted Selective Plane Illumination Microscope (diSPIM) system (Kumar et al. 2014). LSFM has also advanced our understanding *C. elegans* embryogenesis (Chardès et al. 2014; Duncan et al. 2019), such as helping to reveal how the rigid egg shell contributes to asymmetrical cell divisions (Fickentscher and Weiss 2017), how circuit structures are organized within the nerve ring (the *C. elegans* brain) (Moyle et al. 2021), and how the zinc finger protein PIE-1 concentration gradient is established and maintained in the zygote (Benelli et al. 2020).

Although LSFM can also be used to capture embryogenesis in mice (Udan et al. 2014; Ichikawa et al. 2014) and zebrafish (Icha et al. 2016; Kaufmann et al. 2012; Keller et al. 2008; Pang et al. 2020), the increased tissue size and thickness, tissue pigmentation, and lack of transparency limits post-embryonic imaging in these animal models. In contrast, the small size and transparency of *C. elegans* larvae and adults makes them ideal to examine post-embryonic developmental and physiological processes. *C. elegans* is also amenable to high-resolution live imaging of genetically encoded fluorophores fused to proteins to follow protein dynamics and assess gene expression levels and patterns (Keeley et al. 2020; Heppert et al. 2018; Tsuyama et al. 2013; Yoshida et al. 2017; Mita et al. 2019). Genetically encoded fluorophores can also be conjugated to biosensors, which have been used to quantitatively monitor cell cycle state (Adikes et al. 2020) and ATP in *C. elegans* larvae (Garde et al. 2022). *C. elegans* can also be easily stained with vital dyes (Kelley et al. 2019; Schultz and Gumienny 2012; Hermann et al. 2005).

Despite the advantages of LSFM in *C. elegans* for live imaging, LSFM use in larvae and adults has been limited by the difficulty of sample mounting. Low-melt agarose, a common mounting medium used in other model systems, has a gelling temperature of ∼27°C (Icha et al. 2016; Hirsinger and Steventon 2017), which is higher than the upper tolerance of ∼25°C for normal development and physiology of *C. elegans* (Stiernagle 2006). To avoid high temperatures, photo-activated polyethylene glycol (PEG) hydrogels have been used to physically immobilize *C. elegans* for live imaging (Burnett et al. 2018). However, the refractive indices of these hydrogels are often not well-matched for the imaging media or the organism. Here we present a simple protocol for preparing and mounting post-embryonic *C. elegans* for LSFM imaging using a combination of the refractive index matched, ultraviolet (UV)-activated adhesive hydrogel BIO-133 (Han et al. 2021) and FEP tube encasement. We show how this protocol can be used to time-lapse image PVD neuron dendritic branching and pruning. We also demonstrate how this protocol is applicable to imaging a variety of proteins and structures, including extracellular matrix proteins (type IV collagen and laminin), the nuclear envelope, and the distal tip cell (DTC). We expect the adoption of these methods will enable better live-imaging studies of important dynamic cell and developmental processes, such as germ stem cell biology, cell migration, cell division, and cell invasion (Sherwood and Plastino, 2018; Gordon et al., 2020; Smith et al., 2022). Furthermore, this protocol is generalizable and applicable to other organisms with little or no modifications.

## 2. METHODS

### 2.1. Objectives and Validation

Our objective was to develop a procedure for immobilizing larvae and adult *C. elegans* for two-to-three-hour long LSFM timelapse imaging sessions. To accomplish this, we developed a mounting strategy that combines anesthesia, the recently developed BIO-133 UV-activated adhesive hydrogel (Han et al. 2021) and animal encasement in an FEP tube (**Figure 1**). This mounting method allows liquid perfusion of the worms for long term live imaging (upper limit of 3 hours to avoid physiological changes that occur from starvation) and is refractive index-matched to water to minimize the light interface resulting in optimal resolution during imaging. Furthermore, this mounting protocol can be adapted to work with LSFM systems equipped with either universal stage sample mounts (**Figure 1 A-B**) or vertical mounts (**Figure 1 A, C**). To validate our mounting protocol, we used the diSPIM (Kumar et al. 2014) to timelapse image the PVD neurons using a strain harboring endogenously yellow fluorescent protein (YFP) tagged RAB-10 (strain *wy1001[zf1::yfp::rab-10]*) and a membrane tethered GFP expressed in the PVD and OLL neurons (*wyIs592*[*ser-2prom3p::myr-GFP*]). *Rab-10* is a small GTPase involved in post-Golgi vesicle trafficking and is a reporter for the Golgi and early endosome vesicles in the PVD neurons (**Figure 2 A**) (Zou et al. 2015). The multi-dendritic mechanosensory PVD neurons exist as a pair, PVDL and PVDR. Each PVD neuron sits on one side of the animal and has a single axon that extends ventrally to the nerve cord (**Figure 2 A, bottom**). PVD dendritic branching is predictable and developmentally regulated. Specifically, early in the L2 larval stage, the PVD extends 3 processes – one ventrally, one anteriorly, and one posteriorly. Beginning in late L2, the anterior and posterior processes send out short extensions that will elaborate into dendritic trees that compose the non-overlapping, anteroposterior repeating structural units of the PVDs referred to as “menorahs” (**Figure 2 B, top**) (Oren-Suissa et al., 2010). The branches of these menorah structures cover most of the body, except for the neck and head, and are labeled in the proximal-distal and chronological order in which they occur: primary (1°), secondary (2°), tertiary (3°), or quaternary (4°) (**Figure 2 B, bottom**) (Smith et al. 2010). Focusing on the PVDs allowed us to validate the efficacy of this protocol with respect to anterior, midbody, and posterior immobilization as well as imaging clarity throughout LSFM-based live cell imaging. Additionally, PVD development has been the subject of previous confocal-based timelapse studies (Zou et al. 2015) and thus provided us with a point of comparison in the validation of this protocol with respect to stereotyped subcellular dynamics and structural development in a two-to-three-hour timeframe (Wang et al. 2021; Chen and Pan 2021).

**Figure 1.**
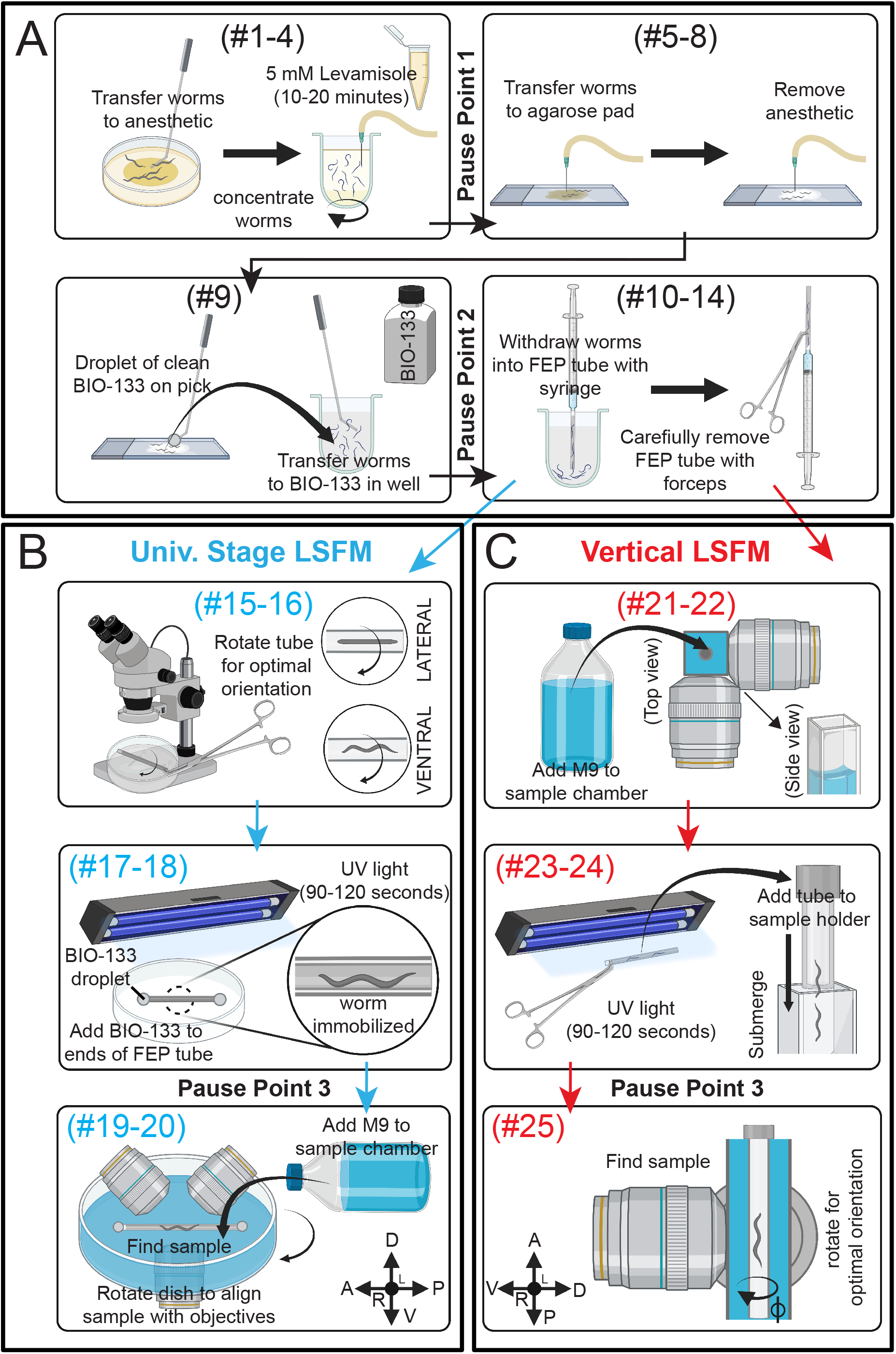
Schematic summary of *C. elegans* post-embryonic BIO-133 mounting strategies for LSFM imaging. **(A)** A schematic summary of steps #1-14 of the FEP-BIO-133 mounting protocol for time-lapse imaging of post-embryonic *C. elegans* on light sheet fluorescence microscopes, including animal anesthesia (top left, *steps #1-4*), transfer to BIO-133 (top right, *steps #5-8*), BIO-133 encapsulation (bottom left, *step #9*), and sample withdrawal into the FEP tube (bottom right, *steps #10-14*). Protocol steps #1-14 can be used for mounting samples on LSFMs configured with either a universal stage mount or a vertically-mounted sample. Pause points #1-2 in the procedure are indicated where they occur in the protocol. **(B)** A schematic summary of FEP tube-sample orientation (top, *steps #15-16*), UV-curation and bonding of FEP tube to Petri dish sample imaging chamber (middle, *steps #17-18*) and sample mounting (bottom, *Steps #19-20*) for LSFM systems equipped with a universal stage mount. After steps #1-14 (A), proceed to steps #15-20. Pause point #3 is indicated. **(C)** A schematic depicting preparation for a vertically-mounted sample, including sample chamber flooding (top, *steps #21-22*), UV-curation and loading of the FEP tube into the sample holder (middle, *steps #23-24*) and rotating the FEP tube to achieve optimal sample orientation (bottom, *step #25*). After steps #1-14 (A), skip steps #15-20 and proceed to steps #21-25. Pause point #3 is indicated.

**Figure 2.**
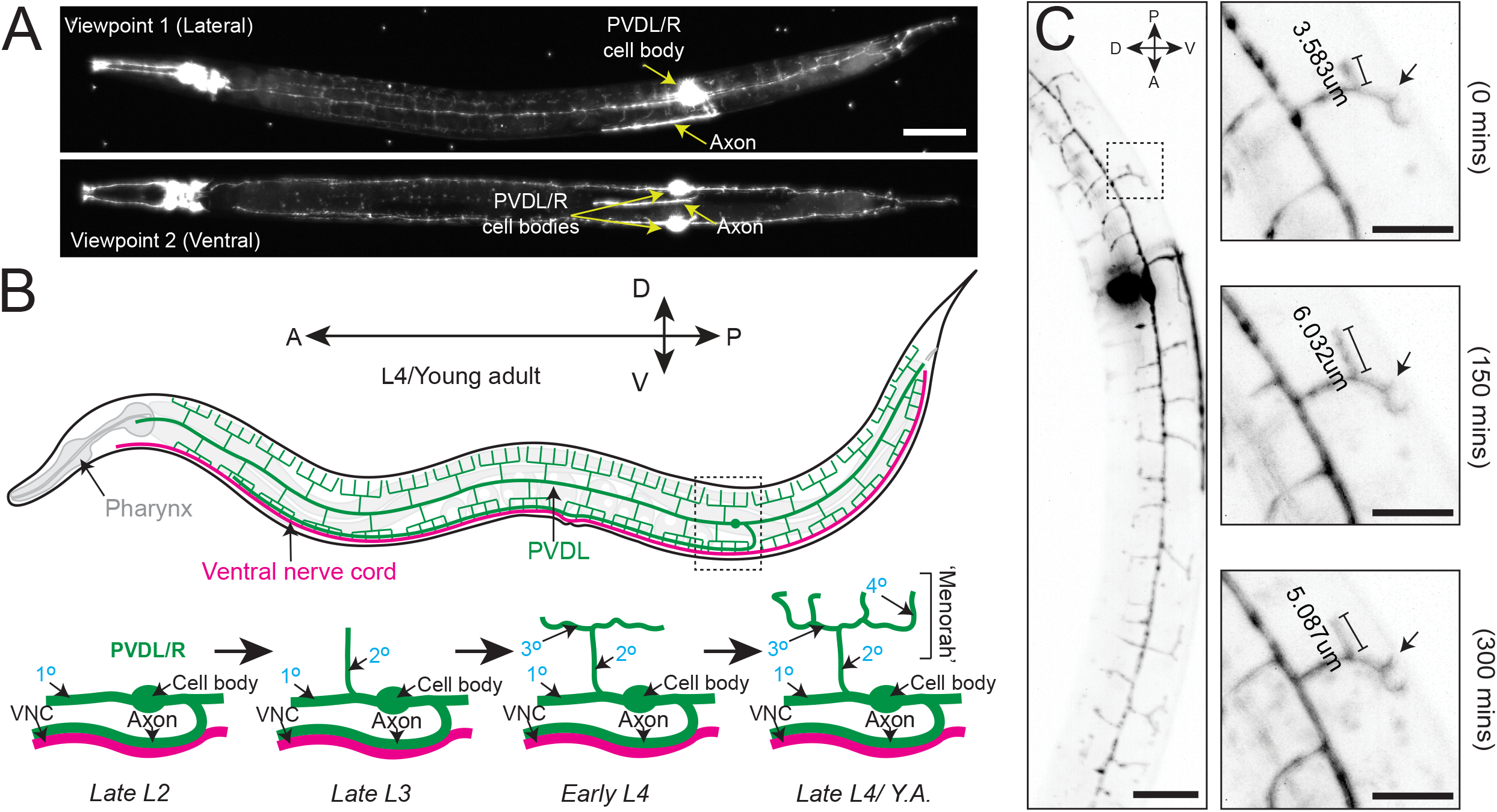
Branching and elongation of PVD neuron dendrites during a 5 hour timelapse on a DiSPIM. **(A)** LSFM Z-projections of an L4 hermaphrodite expressing *yfp::rab-10* (acquired with 40x NA 0.8 water-dipping lenses, z-step = 1 µm) mounted using protocol Steps #1-20 on a diSPIM configured with a universal stage mount. Viewpoints were captured with imaging objectives oriented at 90° to simultaneously view the lateral and ventral aspects of the animal. Scale bar is 25 µm **(B)** (Top) A depiction of the fully elaborated PVD neurons in a young adult hermaphrodite animal. (Bottom) The developmental progression of PVD arborization focusing on the region indicated by the dashed box above. By late L2, the PVD neurons have extended their axons ventrally to contact the nerve cord and the primary (1°) dendrites have elongated along the anterior-posterior axis of the animal. The secondary (2°) dendrites branch dorsally and ventrally from the 1° dendrites by late L3. In early L4, the tertiary (3°) dendrites branch anteroposteriorly from the 2° dendrites, which is followed by the emergence of quaternary (4°) dendrites beginning in the late L4. **(C)** (Left) Timestamp from the beginning of a LSFM timelapse in an L4 hermaphrodite expressing *yfp::rab-10* as in A. (Right) Time series of 3° and 4° dendritic dynamics over the course of a 300 minute LSFM timelapse (acquired with the same parameters described in A). Scale bar 25 µm, 10 µm for inset.

We first performed timelapse imaging of the posterior region of the PVD neuron in an L4 larval stage animal using 2-minute acquisition intervals, a z-step size of 1 µm and z-range of 23 µm (**Movie 1**). This allowed examination of PVD dendritic morphogenesis. We observed tertiary dendritic branch elongation (**Figure 2 C, bracket)** as well as the growth of a quaternary branch (**Figure 2 C, arrow**) (Smith et al. 2010; Albeg et al. 2011).

To further test the compatibility of this mounting protocol with other LSFMs, we imaged multiple fluorescently tagged strains on the Zeiss Lightsheet 7 from two different acquisition angles. Compared to the diSPIM, which is equipped with a universal stage, the Lightsheet 7 has a vertical tube mount, which enables sample rotation during the acquisition for multi-view imaging. Using tiling and a small step size (0.30 µm), we imaged endogenously tagged type IV collagen (EMB-9::mRuby2, **Figure 3 A)**, endogenously tagged laminin (LAM-2::mNG, **Figure 3 B)**, endogenously tagged nucleoporin (NDC-1::mNG, **Figure 3 C)**, and a cell-specific transgene expressing membrane bound GFP in the somatic distal tip cells of the germline (*lag-2p*::GFP, **Figure 3 D)**. Using a 20X, 1.0 NA objective, we observed fine morphological and cellular structures. For example, we resolved the ring of type IV collagen at the edge of the spermatheca in young adult animals (**Figure 3 A’)**, the laminin network surrounding the epithelial cells of the L4 stage spermatheca (**Figure 3 B’)**, the distribution of nucleoporin in L4 stage germ cells (**Figure 3 C’)**, and the elaborations of the distal tip cell in the young adult stage that enwrap the germ stem cell niche (**Figure 3 D’)**. Applying Multiview-registration [Fiji plugin BigStitcher (Hörl et al. 2019)] during image processing, we were also able to create an isotropic image of type IV collagen by combining two different 180° images of the same worm (**Movie 2)**.

**Figure 3.**
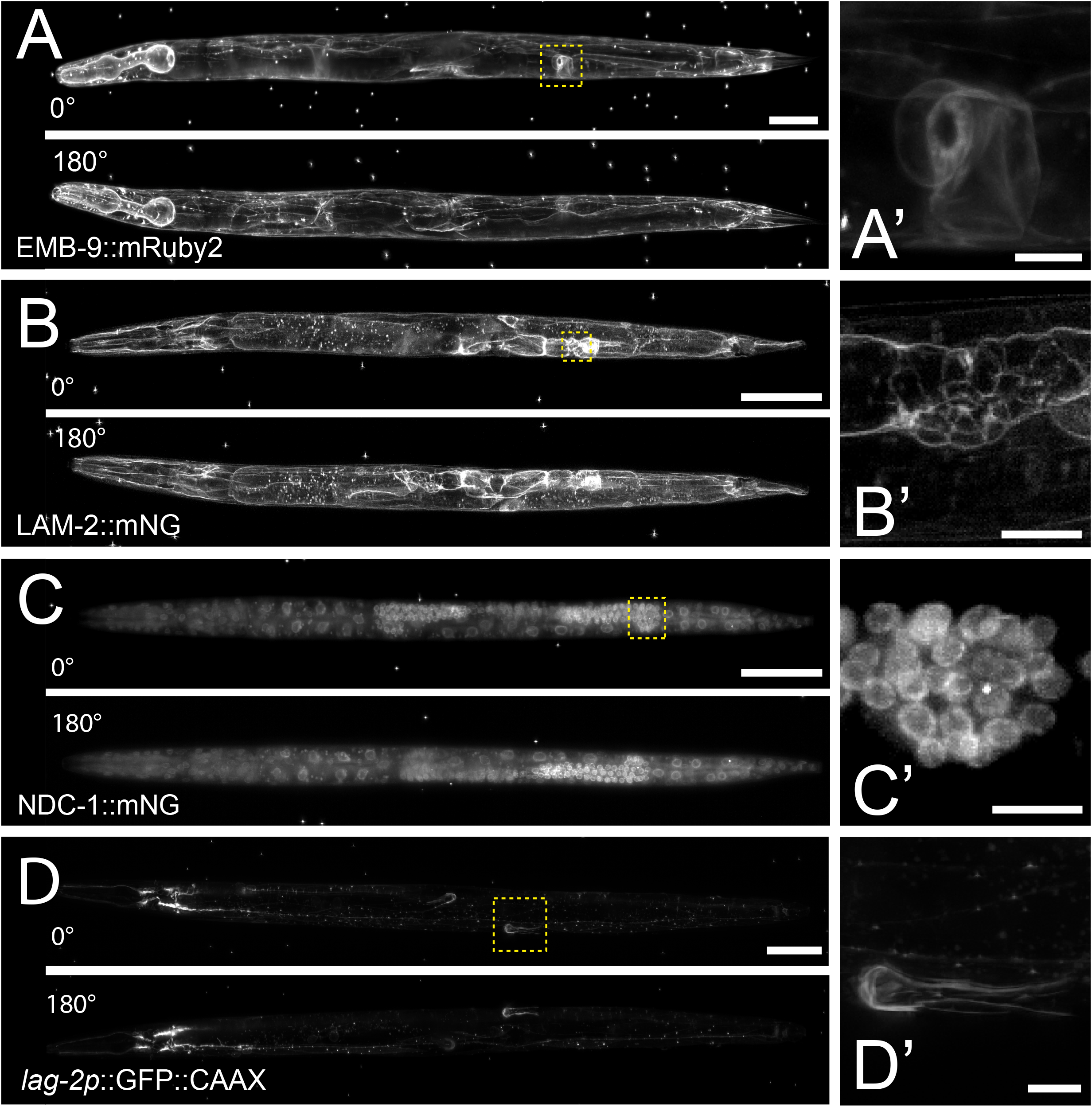
Multiview imaging of endogenously-tagged proteins in *C. elegans* young adults and larvae on a Zeiss Lightsheet 7 with a vertical mount. **(A)** Projected fluorescent images from two viewpoints on the Zeiss L7 showing endogenously-tagged type IV collagen (EMB-9::mRuby2) in a young adult hermaphrodite. The images were acquired from two angles 180° apart using a 20x NA 1.0 water dipping lens (z-step = 0.30 µm). **(B)** Two projected images from LSFM sectioning of endogenously-tagged laminin (LAM-2::mNG) in an L4 hermaphrodite. The images were acquired from two angles 180° apart using a 20x NA 1.0 water dipping lens (z-step = 0.30 µm). **(C)** Two projected images showing endogenously-tagged nucleoporin (NDC-1::mNG) in an L4 hermaphrodite. The images were acquired from two angles 180° apart using a 20x NA 1.0 water dipping lens (z-step = 0.30 µm). **(D)** Two projected images showing distal tip cell (DTC) specific expression of membrane-tethered GFP in an adult hermaphrodite. The images were acquired from two angles 180° apart using a 20x NA 1.0 water dipping lens (z-step = 0.30 µm). Scale bar for all images is 50 µm, 10 µm for inset. **(A’-D’)** Magnified insets of regions in the yellow dashed boxes in A-D.

### 2.2. Materials and Equipment

#### Key Resources

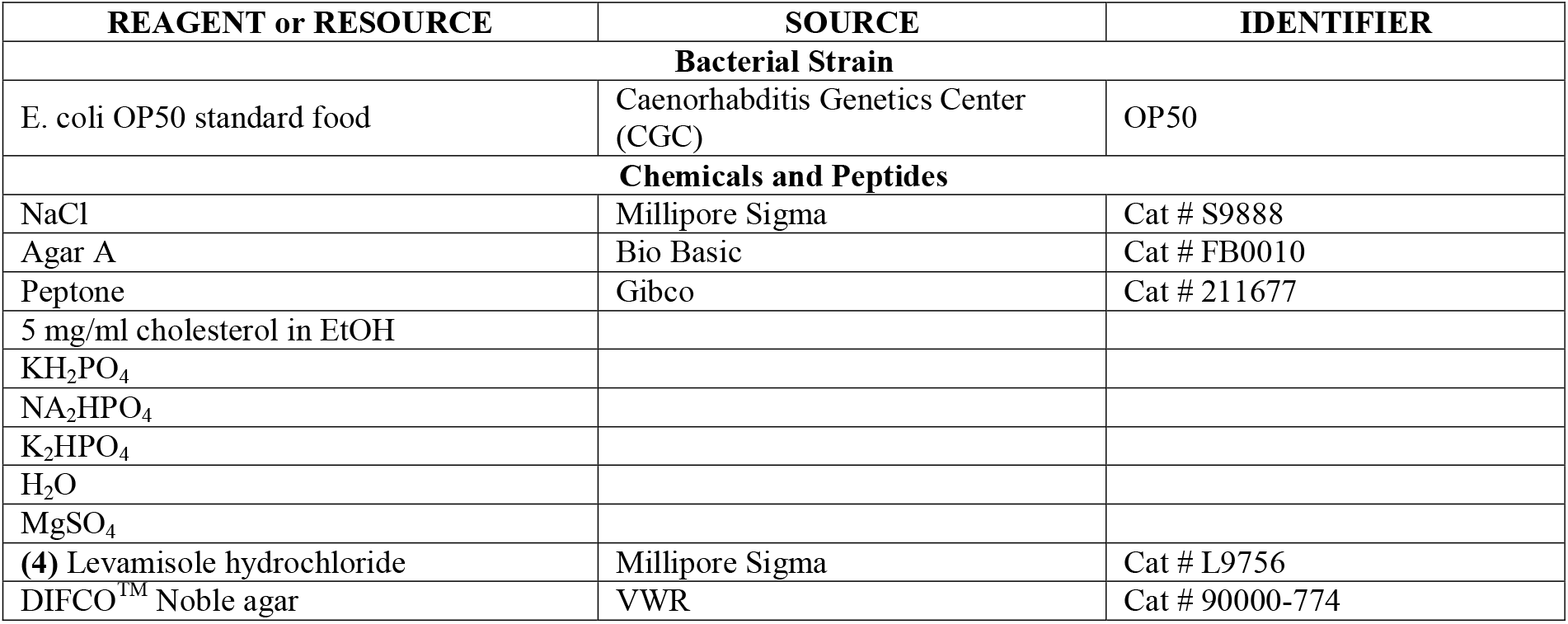
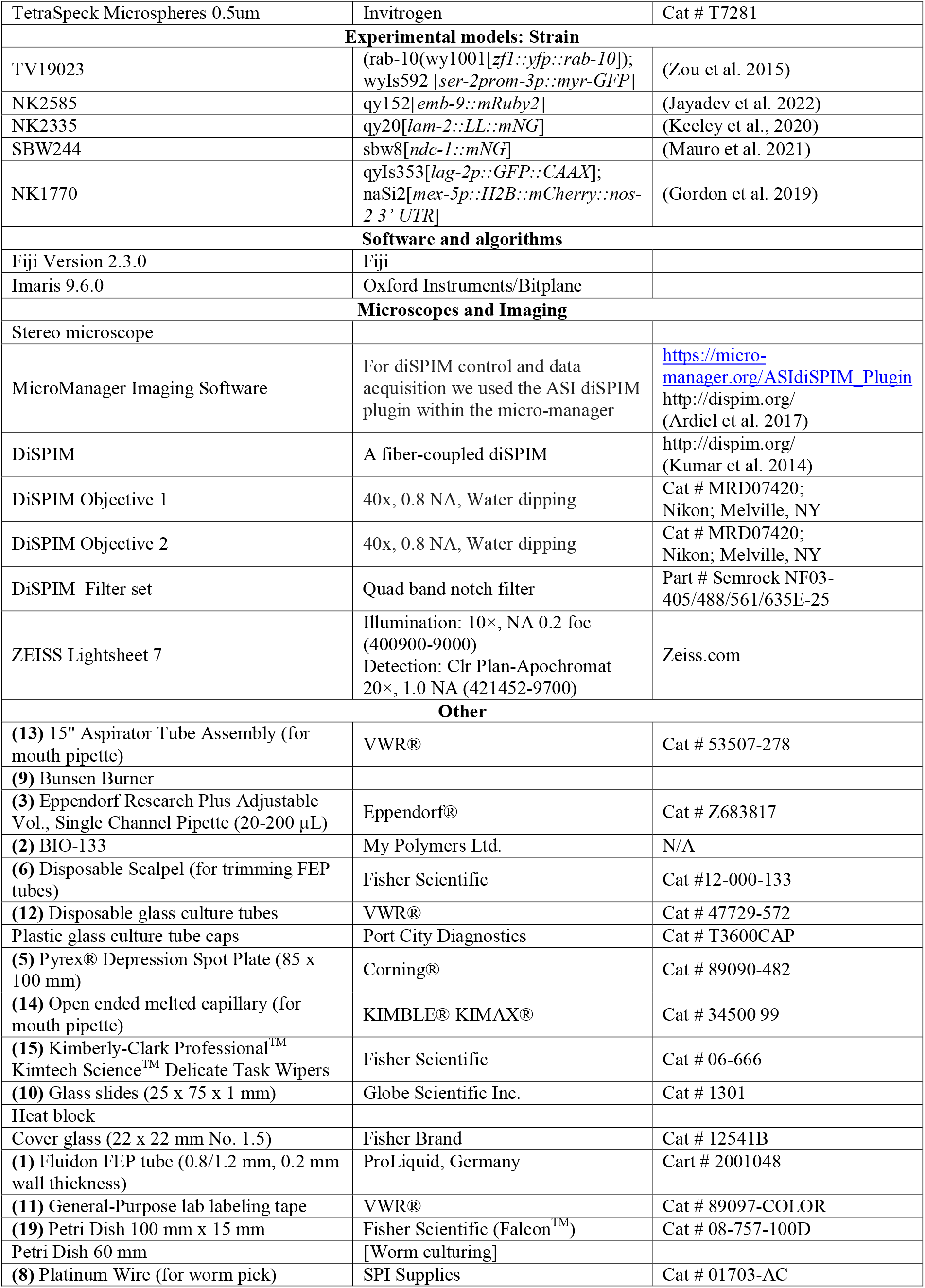
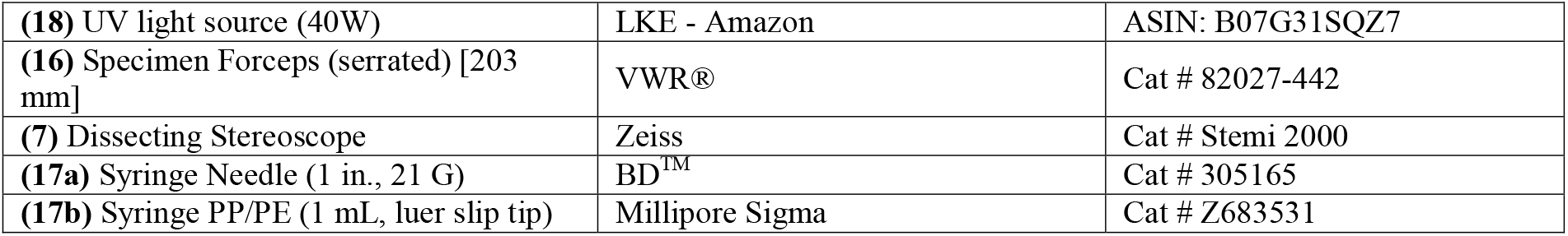

| REAGENT or RESOURCE | SOURCE | IDENTIFIER |
| --- | --- | --- |
| **Bacterial Strain** |  |  |
| E. coli OP50 standard food | Caenorhabditis Genetics Center (CGC) | OP50 |
| **Chemicals and Peptides** |  |  |
| NaCl | Millipore Sigma | Cat # S9888 |
| Agar A | Bio Basic | Cat # FB0010 |
| Peptone | Gibco | Cat # 211677 |
| 5 mg/ml cholesterol in EtOH |  |  |
| KH<sub>2</sub>PO<sub>4</sub> |  |  |
| NA<sub>2</sub>HPO<sub>4</sub> |  |  |
| K<sub>2</sub>HPO<sub>4</sub> |  |  |
| H<sub>2</sub>O |  |  |
| MgSO<sub>4</sub> |  |  |
| (4) Levamisole hydrochloride | Millipore Sigma | Cat # L9756 |
| DIFCO™ Noble agar | VWR | Cat # 90000-774 |
| TetraSpeck Microspheres 0.5um | Invitrogen | Cat # T7281 |
| **Experimental models: Strain** |  |  |
| TV19023 | (rab-10(wy1001[*zfl::yfp::rab-10*]); wyIs592 [*ser-2prom-3p::myr-GFP*]) | (Zou et al. 2015) |
| NK2585 | qy152[*emb-9::mRuby2*] | (Jayadev et al. 2022) |
| NK2335 | qy20[*lam-2::LL::mNG*] | (Keeley et al., 2020) |
| SBW244 | sbw8[*ndc-1::mNG*] | (Mauro et al. 2021) |
| NK1770 | qyIs353[*lag-2p::GFP::CAAX*]; naSi2[*mex-5p::H2B::mCherry::nos-2 3' UTR*] | (Gordon et al. 2019) |
| **Software and algorithms** |  |  |
| Fiji Version 2.3.0 | Fiji |  |
| Imaris 9.6.0 | Oxford Instruments/Bitplane |  |
| **Microscopes and Imaging** |  |  |
| Stereo microscope |  |  |
| MicroManager Imaging Software | For diSPIM control and data acquisition we used the ASI diSPIM plugin within the micro-manager | https://micro-manager.org/ASIdiSPIM_Plugin<br>http://dispim.org/<br>(Ardiel et al. 2017) |
| DiSPIM | A fiber-coupled diSPIM | http://dispim.org/<br>(Kumar et al. 2014) |
| DiSPIM Objective 1 | 40x, 0.8 NA, Water dipping | Cat # MRD07420;<br>Nikon; Melville, NY |
| DiSPIM Objective 2 | 40x, 0.8 NA, Water dipping | Cat # MRD07420;<br>Nikon; Melville, NY |
| DiSPIM Filter set | Quad band notch filter | Part # Semrock NF03-405/488/561/635E-25 |
| ZEISS Lightsheet 7 | Illumination: 10×, NA 0.2 foc (400900-9000)<br>Detection: Clr Plan-Apochromat 20×, 1.0 NA (421452-9700) | Zeiss.com |
| **Other** |  |  |
| (13) 15" Aspirator Tube Assembly (for mouth pipette) | VWR® | Cat # 53507-278 |
| (9) Bunsen Burner |  |  |
| (3) Eppendorf Research Plus Adjustable Vol., Single Channel Pipette (20-200 µL) | Eppendorf® | Cat # Z683817 |
| (2) BIO-133 | My Polymers Ltd. | N/A |
| (6) Disposable Scalpel (for trimming FEP tubes) | Fisher Scientific | Cat #12-000-133 |
| (12) Disposable glass culture tubes | VWR® | Cat # 47729-572 |
| Plastic glass culture tube caps | Port City Diagnostics | Cat # T3600CAP |
| (5) Pyrex® Depression Spot Plate (85 x 100 mm) | Corning® | Cat # 89090-482 |
| (14) Open ended melted capillary (for mouth pipette) | KIMBLE® KIMAX® | Cat # 34500 99 |
| (15) Kimberly-Clark Professional™ Kimtech Science™ Delicate Task Wipers | Fisher Scientific | Cat # 06-666 |
| (10) Glass slides (25 x 75 x 1 mm) | Globe Scientific Inc. | Cat # 1301 |
| Heat block |  |  |
| Cover glass (22 x 22 mm No. 1.5) | Fisher Brand | Cat # 12541B |
| (1) Fluidon FEP tube (0.8/1.2 mm, 0.2 mm wall thickness) | ProLiquid, Germany | Cart # 2001048 |
| (11) General-Purpose lab labeling tape | VWR® | Cat # 89097-COLOR |
| (19) Petri Dish 100 mm x 15 mm | Fisher Scientific (Falcon™) | Cat # 08-757-100D |
| Petri Dish 60 mm | [Worm culturing] |  |
| (8) Platinum Wire (for worm pick) | SPI Supplies | Cat # 01703-AC |
| (18) UV light source (40W) | LKE - Amazon | ASIN: B07G31SQZ7 |
| (16) Specimen Forceps (serrated) [203 mm] | VWR® | Cat # 82027-442 |
| (7) Dissecting Stereoscope | Zeiss | Cat # Stemi 2000 |
| (17a) Syringe Needle (1 in., 21 G) | BD™ | Cat # 305165 |
| (17b) Syringe PP/PE (1 mL, luer slip tip) | Millipore Sigma | Cat # Z683531 |

#### M9 Buffer*

M9 Buffer\*
| REAGENT | FINAL CONCENTRATION | AMOUNT |
| --- | --- | --- |
| Na<sub>2</sub>HPO<sub>4</sub> | 42.2 mM | 6 g |
| KH<sub>2</sub>PO<sub>4</sub> | 22 mM | 3 g |
| NaCl | 85.5 mM | 5 g |
| 1 M MgSO<sub>4</sub> | 1 mM | 1 mL |
| Deionized water | - | 999 mL |
| **Total** | - | **1 L** |

\*Autoclave to sterilize. Aliquot 50 mL into 50 mL falcon tubes. One aliquot will provide enough imaging buffer for 1 timelapse imaging session.

#### Nematode growth medium (NGM) agar plates*

Nematode growth medium (NGM) agar plates\*
| REAGENT | FINAL CONCENTRATION | AMOUNT |
| --- | --- | --- |
| Agar A | 17 g/L | 34 g |
| Peptone | 2.5 g/L | 5 g |
| NaCl | 25.66 mM | 3 g |
| Cholesterol (5 mg/mL) | 12.92 µM | 2 mL |
| Deionized water | - | 1.95 L |
| **Total** | - | **2 L** |

\*Sterilize with autoclave (60 minutes). Cool to 55°C in a water bath and then add 50 mL 1 M KPO<sub>4</sub> buffer (pH 6.0), 2 mL 1 M MgSO<sub>4</sub>, and 2 mL 1 M CaCl<sub>2</sub>. Add 8 mL of warm NGM to each sterile plastic Petri dish using sterile technique and allow to cool. For storage, plates are inverted (NGM side up) at 4°C. NGM plates are warmed to room temperature before seeding with OP50 bacteria for feeding and culturing *C. elegans* strains.

#### Levamisole stock solution (anesthetic)

1. Prepare 200 mM levamisole stock solution in sterile water.
2. Aliquot 150 µL anesthetic stock solution into 1.5 mL Eppendorf tubes and store at -20°C.

#### 4% (weight/volume) noble agar

1. Microwave 4% (weight/volume) noble agar in water to dissolve.
2. Aliquot 1 mL of the melted noble agar into disposable glass tubes and cover with foil or plastic cap. Store at room temperature for up to 3 months.
3. To use, melt noble agar in the glass tube over a Bunsen burner and add to heat block at 70°C to prevent solidification.

### 2.3 Stepwise Procedures

Steps 1-14 described below are shown in **Figure 1 A** and **Supplemental Movie 1**. Video tutorials for agar pad construction, worm anesthetization, and worm transfer can also be found elsewhere (Kelley et al. 2017). All necessary materials required to perform this procedure following preparation of M9 and NGM plates are shown in **Supplemental Figure 1**.

**Movies can be found at: https://doi.org/10.6084/m9.figshare.20443110.v4**

**Total time:** 45-65 minutes

#### *C. elegans* stage selection and anesthesia (Timing: ∼30 minutes)

1. Synchronize worm cultures (Porta-de-la-Riva et al. 2012) (**Time:** 15 minutes) or pick appropriate staged animals for imaging. (**Time:** 2-3 minutes)
2. Add 50 µL anesthesia solution (5 mM Levamisole in M9) to a clean well in a glass depression dish. <u>Alternative to Anesthesia:</u> In addition to immobilization, the anesthetic relaxes the animals into a straight conformation, which facilitates consistent tissue geometry during imaging and permits Multiview registration. However, the use of anesthetic is not suitable for all experiments as levamisole is an acetylcholine receptor agonist that results in muscle contraction (Manjarrez and Mailler 2020). As an alternative, we found animals can be immobilized with cold temperatures by treatment at 5-7°C for ∼15 minutes prior BIO-133 UV-crosslinking.
3. Add 100 µL of BIO-133 to a clean well of the glass depression dish. <u>Detail for precision:</u> BIO-133 is very viscous. Use a scalpel to trim the end of a pipette tip to transfer the hydrogel more easily. *(For Multiview registration)* In an Eppendorf tube combine 80uL BIO-133 and 20 µL of TetraSpeck Microspheres (1:2000 dilution), vortex thoroughly to ensure beads are evenly dispersed in BIO-133. Once mixed, add 50-100 µL of BIO-133 to a clean well.
4. Transfer 20-50 animals to the anesthesia solution and wait for 12 minutes or until most of the animals have ceased moving. Larvae or adults should be straight and rod-like before proceeding to the next step. (**Time**: 15-20 minutes) **<u>Transferring</u> *<u>C</u>***. ***<u>elegans</u>* <u>from anesthetic to BIO-133</u>** (**Timing: ∼15 minutes)**
5. First swirl the glass depression dish to concentrate the anesthetized animals in the center of the well and then use the mouth pipette to remove most of the liquid anesthetic from the well to further concentrate the worm bodies. (**Time:** 1-3 minutes)
6. Prepare an agar pad on a glass slide (See Kelley et al., 2017 for details on agar pad construction) and allow to cool for 1 min. (**Time:** 1-2 minutes)
7. Use a mouth pipette to transfer anesthetized animals from the well in the glass depression dish to the agar pad. (**Time:** 1 minute)
8. Use a mouth pipette to remove anesthetic liquid from the agar pad until animals appear nearly dry (**Supplemental Figure 2**). Avoid removing anesthetized animals with the anesthetic solution. (**Time:** 1-3 minutes)
9. Using a worm pick, gather a droplet of BIO-133 at the end of the pick. Use the BIO-133 droplet to pick and then transfer worms from the nearly dry agar pad to the well of the glass depression dish that contains the BIO-133. Carefully and vigorously swirl the worms in the BIO-133 to separate individual animals and break up any liquid droplets or bubbles that form from the worm transfer. (**Time:** 2-5 minutes) <u>Detail for precision:</u> Any transfer of the anesthetic or water to the BIO-133 solution will result in droplets forming in the adhesive, which will trap the animals, removing them from the hydrogel. <u>Detail for precision:</u> Transferring individual animals rather than many larvae or adults on the pick at the same time will reduce the chances of aggregation. **<u>Loading BIO-133-encased</u> *<u>C</u>***. ***<u>elegans</u>* <u>into FEP tube and polymerizing the mount</u> (Timing: ∼20 minutes)**
10. Attach the 21-G syringe needle to the 1mL syringe barrel.
11. Use serrated forceps to slide the FEP tube onto the 21-G syringe needle. <u>Detail for precision:</u> FEP tubes need to be rinsed and stored in double-distilled water prior to use (reference https://huiskenlab.com/sample-mounting/). Dry the outside of the tube with a Kimwipe and push air through the tube using the syringe plunger to dry the inside of the tube (**Time:** 1-3 minutes). Removing all water will reduce the number of droplets in the BIO-133. <u>Detail for precision:</u> Depending on the length of the FEP tube, it may be necessary to use a disposable scalpel or razor blade to trim the tube into 2-5 cm lengths. Having a shorter segment of FEP tube reduces the time required to find the sample on a LSFM system by minimizing the area containing the sample. Shorter segments of FEP tubes also bond more easily to the bottom of the plastic Petri dish that will become the imaging chamber (See steps 14-16).
12. Place the open end of the FEP tube that is attached to the syringe into the BIO-133 adhesive solution. Using the syringe plunger, draw BIO-133 into the FEP tube until the tube is ¼ full. This primes the tube and ensures that *C. elegans* larvae and adults are positioned centrally, away from the edge of the FEP tube (Step 17). (**Time:** 1-3 minutes) <u>Detail for precision:</u> Due to the high viscosity of the BIO-133 adhesive solution, there will be a delay between when you stop pulling the syringe plunger and when BIO-133 stops flowing into the FEP tube. If more than ¼ of the tube is filled with BIO-133 by the time the pressure is equalized, carefully expel the excess BIO-133 back into the well of the glass depression dish. <u>Detail for precision:</u> To avoid introducing air bubbles into the FEP tube, do not remove the end of the tube from the BIO-133 until you have filled the final ¾ with anesthetized animals and BIO-133 (Step 14).
13. Slowly pull the plunger to draw 5-10 anesthetized animals into the primed FEP tube. (**Time:** 2-5 minutes) <u>Detail for precision:</u> Due to the high viscosity of the BIO-133 adhesive solution, there will be a delay between pulling the syringe plunger and drawing anesthetized animals into the FEP tube. To avoid drawing BIO-133 and animals into the syringe barrel, stop pulling the syringe plunger when the FEP tube is ¾ full. Wait until the pressure equalizes, the FEP tube is full, and the worms stop flowing before removing the end of the FEP tube from the BIO-133 to avoid introducing air bubbles to the FEP tube. <u>Detail for precision:</u> Position the opening of the FEP tube so that the animals will be drawn into the tube longitudinally. Draw one animal up at a time and avoid overlapping animals in the tube. <u>Detail for precision:</u> Ensure that larvae and adults occupy the middle of the FEP tube since LSFM systems equipped with dip lenses will not be able to image animals that are too close to the ends of the FEP tube.
14. Remove the FEP tube from the BIO-133 and check the open end of the FEP tube and the end connected to the needle for air bubbles. The FEP tube should be filled with the adhesive solution, *C. elegans* larvae and adults, and free of air bubbles. Detach the FEP tube from the syringe with serrated forceps. (**Time:** 1-2 minutes) **IF USING A VERTICAL MOUNT, SKIP TO STEPS 21-25** <u>(Steps 15-20 described below are shown in</u> **<u>Figure 1 B</u>** <u>and</u> **<u>Supplemental Movie 2</u>**)
15. Place the FEP tube in the middle of the Petri dish. Add 2-3 drops of BIO-133 hydrogel to the FEP tube using a worm pick or pipette tip. BIO-133 will stabilize the FEP tube during and following UV-treatment. (**Time:** 1-2 minutes)
16. Use a stereo microscope to find the optimal orientation of the FEP tube such that your sample is as close as possible to the imaging objective. If multiple animals are mounted, roll the FEP tube in the uncured BIO-133 to achieve the orientation in which most animals are oriented properly (**Figure 1 B**). (**Time:** 1-3 minutes)
17. Cure the mount with UV light for 2 minutes to crosslink the BIO-133 around the anesthetized animals and bond the sample-containing FEP tube to the plastic Petri dish imaging chamber. (**Time:** 2 minutes) **<u>Installing the mount on an LSFM equipped with a universal stage and dipping lenses (Timing: ~2 minutes)</u>**
18. After UV curing, the FEP tube should be stably attached to the surface of the plastic Petri dish and the sample should be encased in a rigid hydrogel in the FEP tube. Ensure that the FEP tube is securely attached to the Petri dish by lightly tapping it with forceps or a pipette tip. The tube should not budge or move at all before proceeding. (**Time:** 1 minute)
19. Add mount to the universal stage on the LSFM system. Once the mount is resting on the universal stage, rotate the dish until your sample is optimally aligned with the imaging objectives (**Figure 1 B**). Fasten the specimen clips to secure the Petri dish imaging chamber. (**Time:** 1 minute)
20. Slowly fill the Petri dish imaging chamber with 45-50 mL room temperature M9 buffer (imaging medium), after which the dipping lens objectives can be lowered into the M9 for sample finding and subsequent imaging. **END OF PROCEDURE FOR LSFM WITH UNIVERSAL STAGE MOUNT** **<u>Installing the mount on an LSFM which requires a vertically mounted sample</u> (Timing: ∼5 minutes)** **<u>(Steps 21-25 described below are shown in Figure 1C and step 23 (UV-curing) is shown in Supplemental Movie 3)</u>**
21. Fill the LSFM media chamber with M9. (**Time:** 1 minute) <u>Detail for Precision:</u> M9 can be added to the media chamber prior to starting the protocol and does not need to be replaced between samples.
22. Wipe the FEP tube containing animals in BIO-133 with a Kimwipe to remove any BIO-133 from the outside of the tube. (**Time:** 1 minute) <u>Detail for Precision:</u> When possible, use forceps to handle the tube to keep the tube as clean as possible, as any smudges on the outside of the tube might impede the clarity of the imaging
23. Cure the mount with UV light for 2 minutes to crosslink the BIO-133 around the anesthetized animals; this can be done before or after detaching the FEP tube from the syringe needle. (**Time:** 2 minutes) <u>Detail for Precision:</u> Use a stereomicroscope to locate the straight, centered, and non-overlapping animals within the FEP tube. (**Time:** 1 minute)
24. Attach the tube in the sample holder, keeping in mind the positions of the animals as identified in step 24. If the animals are close to the end of the tube, place the opposite end of the tube in the sample holder. (**Time:** 1 minute)
25. Place the sample holder with FEP tube back into the mount so that the FEP tube is submerged in M9 and ready for sample finding and imaging. (**Time:** 1 minute)

## 3. ANTICIPATED RESULTS

This work introduces the advantages of LSFM live imaging to long term postembryonic *C. elegans* development, including faster acquisition speed and reduced phototoxicity and photobleaching. Prior to the development of this protocol, light-sheeting imaging of *C. elegans* had been limited to embryos, very short timelapse imaging of larvae and adults, and fixed samples (Chardès et al. 2014; Duncan et al. 2019; Chen et al. 2014; Breimann et al. 2019; Liu et al. 2018). We anticipate that adult or larval encasement in BIO-133 within an FEP tube will enable continuous LSFM imaging for at least 2 hours, a time span that is comparable to that typical of confocal timelapses (Kelley et al. 2017) and which approaches the physiological limit imposed by starvation (Schindler and Sherwood 2014). Unlike the confocal time-lapse mount, this protocol exposes animals to minimal amounts (up to 2 mins) of direct UV light or low temperatures (7°C for the thermal immobilization method). We expect this protocol will thus allow investigations into DNA damage, UV-induced stress, or thermal hyperalgesia (Deng et al. 2020; Plagens et al. 2021; Ma and Shen 2012).

A major advantage of this procedure is low material cost and accessibility of reagents and equipment (See Materials and Equipment table). The mounting strategy can be easily performed with resources already present in most *C. elegans* labs, except for BIO-133 and FEP tubes. Although we only used plastic dishes in the development of this protocol, BIO-133 can be used to bond FEP tubes to glass Petri dishes for a reusable sample chamber.

Compared to the short amount of time between preparing a traditional timelapse slide and imaging a sample on a point-scanning confocal system (Kelley et al. 2017), a limitation of this protocol is the length of time it takes to compose and cure the mount (∼30 minutes) before imaging. In this protocol animals are removed from food for a longer period before imaging, which reduces the time available for timelapse before starvation by ∼30 minutes compared to a slide-based timelapse mount (Kelley et al. 2017). Additionally, since the orientation of animals within the FEP tube is fixed after UV curing, it can take multiple mounting attempts to achieve optimal animal orientation. This protocol is therefore comparatively low throughput. This is a significant drawback to the investigation of developmental processes with sensitive timing, or if there is limited time available to use an LSFM system. To shorten the time to imaging, multiple LSFM timelapse mounts can be assembled in parallel.

Finally, we have not tested the diffusion mechanics of the activated BIO-133 hydrogel. It is possible that this protocol cannot be adapted for use in combination with diffusible cues and hormones (e.g., auxin for degron-based protein depletion) (Zhang et al. 2015; Martinez and Matus 2020; Martinez et al. 2020) or mitogens (Monsalve et al. 2019). However, pre-treatment with drugs or hormones prior to mounting animals may be sufficient to capture the desired effects, depending on the mechanics of the biological process or technique of interest. Since the ends of the FEP tubes are left open in the mount, the BIO-133 hydrogel matrix and sample should also be exposed to oxygen and media.

## 4. DISCUSSION

Here we describe a simple protocol for collecting high-quality post-embryonic LSFM timelapse imaging data of larval and adult *C. elegans*. It is likely that this protocol can be adapted for the purposes of imaging other animal models, as the BIO-133 adhesive is biocompatible and FEP tubes are available in a variety of lengths and diameters. Though this method of immobilization and sample mounting provides novel opportunities for *in vivo* imaging of post-embryonic *C. elegans*, such as germ cell divisions, DTC migrations, sex myoblast migration, and anchor cell invasion, there remain a few shortcomings, such as the extended time it takes to prepare samples as discussed in the anticipated results section (Gordon et al. 2020; Sherwood and Plastino 2018; Adikes et al. 2020).

Among the many advantages to light-sheet microscopy mentioned above, this protocol enables multi-view image data via multidirectional illumination or sample rotation by providing access to the input image data necessary for 4D image reconstruction (Huisken and Stainier 2009; Schmid and Huisken 2015). Using 4D image reconstruction, we were able to discern the ring structure of type IV collagen in the spermathecal valve that opens to the uterus and laminin tightly covering the individual epithelial cells of the spermatheca. The BIO-133 can also be seeded with fluorescent beads (microspheres) as fiduciary markers (Wu et al. 2013; Preibisch et al. 2010) to improve multi-view image processing with greater temporal and spatial registration (**Movie 2)**. This protocol for *C. elegans* post-embryonic timelapse imaging should be adaptable to any light sheet or confocal microscope that contains water dipping lenses and a universal stage mount or vertically mounted samples submerged in a sample chamber.

## 5. FIGURE LEGENDS

**Movie 1. Elaboration of the PVD neuron in the L4 midbody**. A 5 hour timelapse of an L4 hermaphrodite expressing *yfp::rab-10*. The timelapse was acquired on diSPIM with 40x NA 0.8 water-dipping objective lenses and images were collected every 2 minutes.

**Movie 2. Using microspheres for enhanced spatiotemporal resolution**. An isotropic image of endogenously-tagged type IV collagen (EMB-9::mRuby2) derived from the Multiview registration of images in **Figure 3A**.

**Supplemental Movie 1. Protocol Instructional Video 1** – *Step 1 to step 14*

**Supplemental Movie 2. Protocol Instructional Video 2** – *Step 15 to step 17*

**Supplemental Movie 3. Protocol Instructional Video 3** – *Step 23 (UV-curing mount for LSFMs which require vertically-mounted sample)*

## Author Contributions

J.J.S., I.W.K, and D.Q.M conceptualized the project. J.J.S and I.W.K designed the protocol, collected all data (with microscopy and image processing help from C.W. and A.K.), and wrote the manuscript. D.Q.M, D.R.S edited and revised the manuscript. C.W. and A.K. provided additional comments on the manuscript. R.C. independently tested the protocol and provided helpful feedback.

## Acknowledgments

The authors would like to thank J. Henry for help recording the videos and the Marine Biological Laboratory (MBL), the MBL Central Microscopy Facility, and the MBL Embryology course directors, staff, and students for providing the lab space and environment within which this protocol was developed. Michael Weber from the Flamingo team is greatly acknowledged for providing FEP tubes and support during preliminary imaging experiments.

## Bibliography

1. Adikes, R.C., Kohrman, A.Q., Martinez, M.A.Q., et al. 2020. Visualizing the metazoan proliferation-quiescence decision in vivo. eLife 9.

2. Albeg, A., Smith, C.J., Chatzigeorgiou, M., et al. 2011. C. elegans multi-dendritic sensory neurons: morphology and function. Molecular and Cellular Neurosciences 46(1), pp. 308–317.

3. Albert-Smet, I., Marcos-Vidal, A., Vaquero, J.J., Desco, M., Muñoz-Barrutia, A. and Ripoll, J. 2019. Applications of Light-Sheet Microscopy in Microdevices. Frontiers in Neuroanatomy 13, p. 1.

4. Ardiel, E.L., Kumar, A., Marbach, J., et al. 2017. Visualizing calcium flux in freely moving nematode embryos. Biophysical Journal 112(9), pp. 1975–1983.

5. Benelli, R., Struntz, P., Hofmann, D. and Weiss, M. 2020. Quantifying spatiotemporal gradient formation in early Caenorhabditis elegans embryos with lightsheet microscopy. Journal of physics D: Applied physics 53(29), p. 295401.

6. Breimann, L., Preusser, F. and Preibisch, S. 2019. Light-microscopy methods in C. elegans research. Current Opinion in Systems Biology 13, pp. 82–92.

7. Burnett, K., Edsinger, E. and Albrecht, D.R. 2018. Rapid and gentle hydrogel encapsulation of living organisms enables long-term microscopy over multiple hours. Communications Biology 1, p. 73.

8. Chardès, C., Mélénec, P., Bertrand, V. and Lenne, P.-F. 2014. Setting up a simple light sheet microscope for in toto imaging of C. elegans development. Journal of Visualized Experiments (87).

9. Chen, B.-C., Legant, W.R., Wang, K., et al. 2014. Lattice light-sheet microscopy: imaging molecules to embryos at high spatiotemporal resolution. Science 346(6208), p. 1257998.

10. Chen, C.-H. and Pan, C.-L. 2021. Live-cell imaging of PVD dendritic growth cone in post-embryonic C. elegans. STAR Protocols 2(2), p. 100402.

11. Deng, J., Bai, X., Tang, H. and Pang, S. 2020. DNA damage promotes ER stress resistance through elevation of unsaturated phosphatidylcholine in C. elegans. The Journal of Biological Chemistry.

12. Duncan, L.H., Moyle, M.W., Shao, L., et al. 2019. Isotropic Light-Sheet Microscopy and Automated Cell Lineage Analyses to Catalogue Caenorhabditis elegans Embryogenesis with Subcellular Resolution. Journal of Visualized Experiments (148).

13. Fickentscher, R. and Weiss, M. 2017. Physical determinants of asymmetric cell divisions in the early development of Caenorhabditis elegans. Scientific Reports 7(1), p. 9369.

14. Fischer, R.S., Wu, Y., Kanchanawong, P., Shroff, H. and Waterman, C.M. 2011. Microscopy in 3D: a biologist’s toolbox. Trends in Cell Biology 21(12), pp. 682–691.

15. Garde, A., Kenny, I.W., Kelley, L.C., et al. 2022. Localized glucose import, glycolytic processing, and mitochondria generate a focused ATP burst to power basement-membrane invasion. Developmental Cell 57(6), p. 732–749.e7.

16. Girstmair, J., Zakrzewski, A., Lapraz, F., et al. 2016. Light-sheet microscopy for everyone? Experience of building an OpenSPIM to study flatworm development. BMC Developmental Biology 16(1), p. 22.

17. Gordon, K.L., Payne, S.G., Linden-High, L.M., et al. 2019. Ectopic Germ Cells Can Induce Niche-like Enwrapment by Neighboring Body Wall Muscle. Current Biology 29(5), p. 823–833.e5.

18. Gordon, K.L., Zussman, J.W., Li, X., Miller, C. and Sherwood, D.R. 2020. Stem cell niche exit in C. elegans via orientation and segregation of daughter cells by a cryptic cell outside the niche. eLife 9.

19. Han, X., Su, Y., White, H., et al. 2021. A polymer index-matched to water enables diverse applications in fluorescence microscopy. Lab on A Chip 21(8), pp. 1549–1562.

20. Heppert, J.K., Pani, A.M., Roberts, A.M., Dickinson, D.J. and Goldstein, B. 2018. A CRISPR Tagging-Based Screen Reveals Localized Players in Wnt-Directed Asymmetric Cell Division. Genetics 208(3), pp. 1147–1164.

21. Hermann, G.J., Schroeder, L.K., Hieb, C.A., et al. 2005. Genetic analysis of lysosomal trafficking in Caenorhabditis elegans. Molecular Biology of the Cell 16(7), pp. 3273–3288.

22. Hirsinger, E. and Steventon, B. 2017. A versatile mounting method for long term imaging of zebrafish development. Journal of Visualized Experiments (119).

23. Hörl, D., Rojas Rusak, F., Preusser, F., et al. 2019. BigStitcher: reconstructing high-resolution image datasets of cleared and expanded samples. Nature Methods 16(9), pp. 870–874.

24. Huisken, J. and Stainier, D.Y.R. 2009. Selective plane illumination microscopy techniques in developmental biology. Development 136(12), pp. 1963–1975.

25. Icha, J., Schmied, C., Sidhaye, J., Tomancak, P., Preibisch, S. and Norden, C. 2016. Using light sheet fluorescence microscopy to image zebrafish eye development. Journal of Visualized Experiments (110), p. e53966.

26. Ichikawa, T., Nakazato, K., Keller, P.J., et al. 2014. Live imaging and quantitative analysis of gastrulation in mouse embryos using light-sheet microscopy and 3D tracking tools. Nature Protocols 9(3), pp. 575–585.

27. Jayadev, R., Morais, M.R.P.T., Ellingford, J.M., et al. 2022. A basement membrane discovery pipeline uncovers network complexity, regulators, and human disease associations. Science Advances 8(20), p. eabn2265.

28. Kaufmann, A., Mickoleit, M., Weber, M. and Huisken, J. 2012. Multilayer mounting enables long-term imaging of zebrafish development in a light sheet microscope. Development 139(17), pp. 3242–3247.

29. Keeley, D.P., Hastie, E., Jayadev, R., et al. 2020. Comprehensive Endogenous Tagging of Basement Membrane Components Reveals Dynamic Movement within the Matrix Scaffolding. Developmental Cell 54(1), p. 60–74.e7.

30. Keller, P.J., Schmidt, A.D., Wittbrodt, J. and Stelzer, E.H.K. 2008. Reconstruction of zebrafish early embryonic development by scanned light sheet microscopy. Science 322(5904), pp. 1065–1069.

31. Kelley, L.C., Chi, Q., Cáceres, R., et al. 2019. Adaptive F-Actin Polymerization and Localized ATP Production Drive Basement Membrane Invasion in the Absence of MMPs. Developmental Cell 48(3), p. 313–328.e8.

32. Kelley, L.C., Wang, Z., Hagedorn, E.J., et al. 2017. Live-cell confocal microscopy and quantitative 4D image analysis of anchor-cell invasion through the basement membrane in Caenorhabditis elegans. Nature Protocols 12(10), pp. 2081–2096.

33. Kumar, A., Wu, Y., Christensen, R., et al. 2014. Dual-view plane illumination microscopy for rapid and spatially isotropic imaging. Nature Protocols 9(11), pp. 2555–2573.

34. Liu, T.-L., Upadhyayula, S., Milkie, D.E., et al. 2018. Observing the cell in its native state: Imaging subcellular dynamics in multicellular organisms. Science 360(6386).

35. Ma, X. and Shen, Y. 2012. Structural basis for degeneracy among thermosensory neurons in Caenorhabditis elegans. The Journal of Neuroscience 32(1), pp. 1–3.

36. Manjarrez, J.R. and Mailler, R. 2020. Stress and timing associated with Caenorhabditis elegans immobilization methods. Heliyon 6(7), p. e04263.

37. Martinez, M.A.Q., Kinney, B.A., Medwig-Kinney, T.N., et al. 2020. Rapid Degradation of Caenorhabditis elegans Proteins at Single-Cell Resolution with a Synthetic Auxin. G3 (Bethesda, Md.) 10(1), pp. 267–280.

38. Martinez, M.A.Q. and Matus, D.Q. 2020. Auxin-mediated Protein Degradation in Caenorhabditis elegans. Bio-protocol 10(8).

39. Mauro, M.S., Celma, G., Zimyanin, V., Gibson, K.H., Redemann, S. and Bahmanyar, S. 2021. NDC1 is necessary for the stable assembly of the nuclear pore scaffold to establish nuclear transport in early C. elegans embryos. BioRxiv.

40. Mita, M., Ito, M., Harada, K., et al. 2019. Green Fluorescent Protein-Based Glucose Indicators Report Glucose Dynamics in Living Cells. Analytical Chemistry 91(7), pp. 4821–4830.

41. Monsalve, G.C., Yamamoto, K.R. and Ward, J.D. 2019. A New Tool for Inducible Gene Expression in Caenorhabditis elegans. Genetics 211(2), pp. 419–430.

42. Moyle, M.W., Barnes, K.M., Kuchroo, M., et al. 2021. Structural and developmental principles of neuropil assembly in C. elegans. Nature 591(7848), pp. 99–104.

43. Pang, M., Bai, L., Zong, W., et al. 2020. Light-sheet fluorescence imaging charts the gastrula origin of vascular endothelial cells in early zebrafish embryos. Cell discovery 6, p. 74.

44. Plagens, R.N., Mossiah, I., Kim Guisbert, K.S. and Guisbert, E. 2021. Chronic temperature stress inhibits reproduction and disrupts endocytosis via chaperone titration in Caenorhabditis elegans. BMC Biology 19(1), p. 75.

45. Porta-de-la-Riva, M., Fontrodona, L., Villanueva, A. and Cerón, J. 2012. Basic Caenorhabditis elegans methods: synchronization and observation. Journal of Visualized Experiments (64), p. e4019.

46. Preibisch, S., Saalfeld, S., Schindelin, J. and Tomancak, P. 2010. Software for bead-based registration of selective plane illumination microscopy data. Nature Methods 7(6), pp. 418–419.

47. Schindler, A.J. and Sherwood, D.R. 2014. Should I stay or should I go? Identification of novel nutritionally regulated developmental checkpoints in C. elegans. Worm 3(4), p. e979658.

48. Schmid, B. and Huisken, J. 2015. Real-time multi-view deconvolution. Bioinformatics 31(20), pp. 3398–3400.

49. Schultz, R.D. and Gumienny, T.L. 2012. Visualization of Caenorhabditis elegans cuticular structures using the lipophilic vital dye DiI. Journal of Visualized Experiments (59), p. e3362.

50. Sherwood, D.R. and Plastino, J. 2018. Invading, Leading and Navigating Cells in Caenorhabditis elegans: Insights into Cell Movement in Vivo. Genetics 208(1), pp. 53–78.

51. Smith, C.J., Watson, J.D., Spencer, W.C., et al. 2010. Time-lapse imaging and cell-specific expression profiling reveal dynamic branching and molecular determinants of a multi-dendritic nociceptor in C. elegans. Developmental Biology 345(1), pp. 18–33.

52. Smith, J.J., Xiao, Y., Parsan, N., et al. 2022. The SWI/SNF chromatin remodeling assemblies BAF and PBAF differentially regulate cell cycle exit and cellular invasion in vivo. PLoS Genetics 18(1), p. e1009981.

53. Steuwe, C., Vaeyens, M.-M., Jorge-Peñas, A., et al. 2020. Fast quantitative time lapse displacement imaging of endothelial cell invasion. Plos One 15(1), p. e0227286.

54. Stiernagle, T. 2006. Maintenance of C. elegans. Wormbook: the Online Review of C. Elegans Biology, pp. 1–11.

55. Tsuyama, T., Kishikawa, J., Han, Y.-W., et al. 2013. In vivo fluorescent adenosine 5’-triphosphate (ATP) imaging of Drosophila melanogaster and Caenorhabditis elegans by using a genetically encoded fluorescent ATP biosensor optimized for low temperatures. Analytical Chemistry 85(16), pp. 7889–7896.

56. Udan, R.S., Piazza, V.G., Hsu, C.-W., Hadjantonakis, A.-K. and Dickinson, M.E. 2014. Quantitative imaging of cell dynamics in mouse embryos using light-sheet microscopy. Development 141(22), pp. 4406–4414.

57. Wang, X., Li, T., Hu, J., et al. 2021. In vivo imaging of a PVD neuron in Caenorhabditis elegans. STAR Protocols 2(1), p. 100309.

58. Wu, Y., Wawrzusin, P., Senseney, J., et al. 2013. Spatially isotropic four-dimensional imaging with dual-view plane illumination microscopy. Nature Biotechnology 31(11), pp. 1032–1038.

59. Yoshida, T., Alfaqaan, S., Sasaoka, N. and Imamura, H. 2017. Application of FRET-Based Biosensor “ATeam” for Visualization of ATP Levels in the Mitochondrial Matrix of Living Mammalian Cells. Methods in Molecular Biology 1567, pp. 231–243.

60. Zhang, L., Ward, J.D., Cheng, Z. and Dernburg, A.F. 2015. The auxin-inducible degradation (AID) system enables versatile conditional protein depletion in C. elegans. Development 142(24), pp. 4374–4384.

61. Zou, W., Yadav, S., DeVault, L., Nung Jan, Y. and Sherwood, D.R. 2015. RAB-10-Dependent Membrane Transport Is Required for Dendrite Arborization. PLoS Genetics 11(9), p. e1005484.

