## Supplemental Figures and Legends for "A light sheet fluorescence microscopy protocol for *Caenorhabditis elegans* larvae and adults"

Figure S1

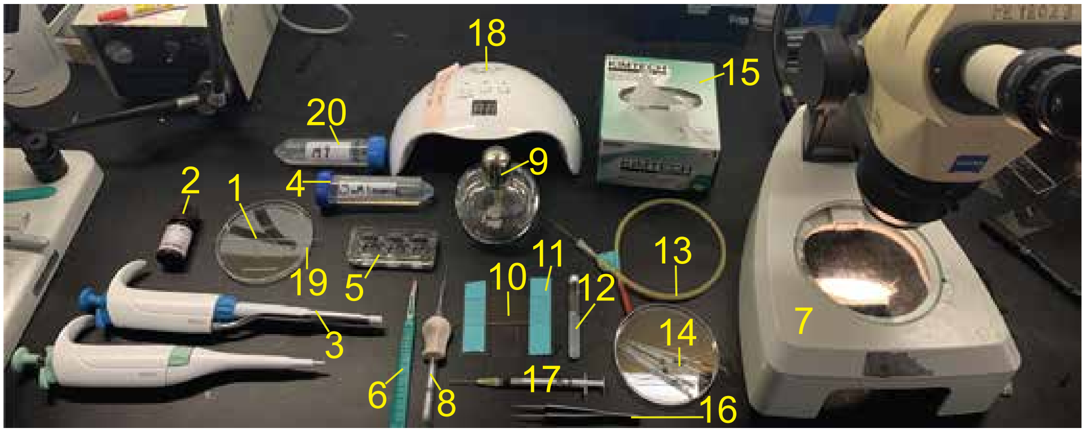


**Supplemental Figure 1. Necessary materials for BIO-133 immobilization protocol.** An image showing all of the materials and tools used for the BIO-133 immobilization protocol. The yellow item numbers label individual materials and tools included in the Materials & Reagents section and correspond to numbers in parentheses preceding each item in the reagents table.

Figure S2

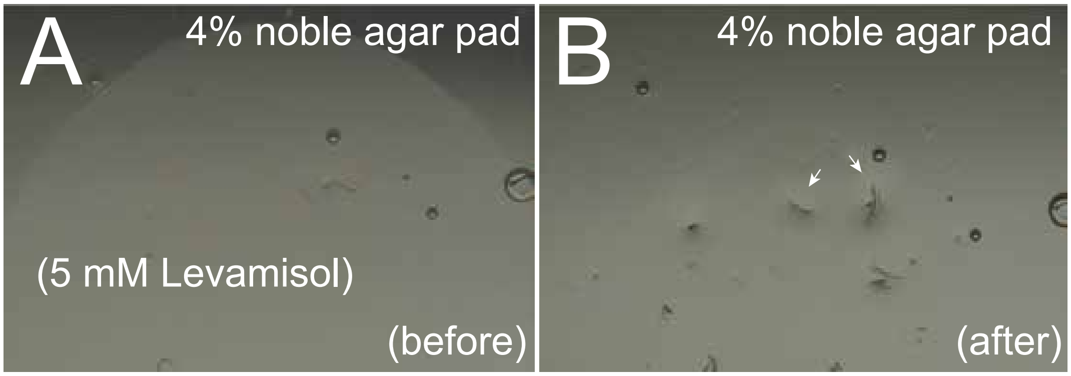


**Supplemental Figure 2. Removing excess levamisole from agar pad for BIO-133 transfer. (A)** A brightfield image through a dissecting microscope shows anesthetized animals in 5mM levamisole on a 4% Noble agar pad after 10 minute anesthesia. **(B)** A brightfield image showing *C. elegans* (arrows) following removal of most of the anesthetic. Before transfer of larvae or adults to BIO-133, a minimal amount of anesthetic is left such that individual animals or groups of animals are collected in small droplets of levamisole.

Figure S3

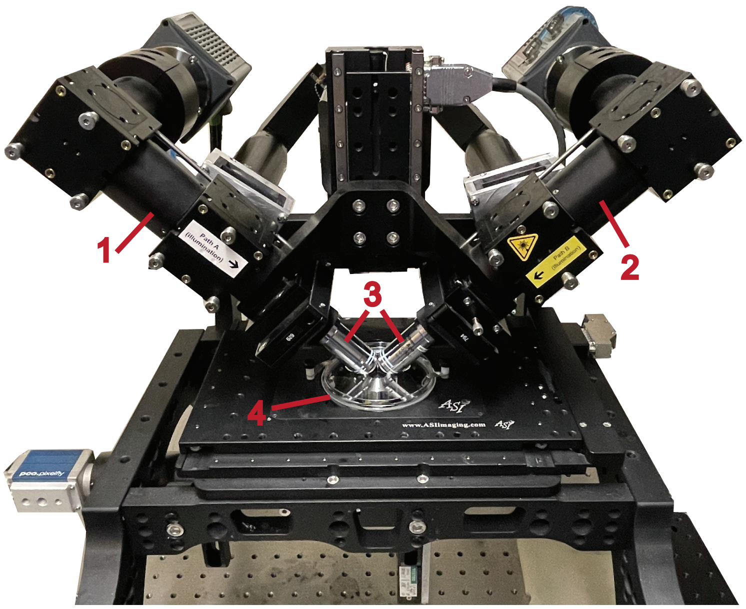


**Supplemental Figure 3. Sample mounted on a diSPIM configured with a universal stage mount (A)** An image of an FEP tube-BIO-133 sample mounted on a diSPIM, which was used to acquire universal stage mounted data presented here. The illumination/detection paths (**1** and **2**), 40X 0.8 NA water dipping lenses (**3**), and the universal stage mount with mounted sample (**4**) are labeled.
